# Diverse Viral Pathogens in Australian Canines: Limited Geographic Structure and the First Detection of an RNA Virus in Dingoes

**DOI:** 10.1101/2025.02.16.638538

**Authors:** Jonathon C.O. Mifsud, Erin Harvey, Kate Van Brussel, Annabelle Olsson, Benjamin J. Pitcher, Jane Hall, Heather Fenton, Brendan F. Alting, Sabrina Sadiq, Edward C. Holmes

**Affiliations:** School of Medical Sciences, The University of Sydney, Sydney, New South Wales, 2006, Australia; Wildlife Health and Conservation Hospital, Sydney School of Veterinary Science, The University of Sydney, Sydney, New South Wales, 2006, Australia; Taronga Institute of Science and Learning, Taronga Conservation Society, Dubbo and Sydney, New South Wales, Australia; School of Natural Sciences, Faculty of Science and Engineering, Macquarie University, Sydney, New South Wales, Australia; Australian Registry of Wildlife Health, Taronga Conservation Society Australia, Mosman, New South Wales, 2088, Australia; Centre for Ecosystem Science, School of Biological, Earth and Environmental Sciences, University of New South Wales, Sydney, New South Wales, Australia

**Keywords:** domestic dog, dingo, Australia, virus discovery, phylogeny, metatranscriptomics

## Abstract

Viruses impose a substantial disease burden on dogs and the close relationship between dogs and humans may facilitate zoonotic disease emergence. Australia’s geographic isolation, strict biosecurity measures and native dingo populations present a unique model for understanding the spread and evolution of canine viruses. However, aside from a few well-characterised pathogens, genomic data are scarce for many common dog viruses, limiting our understanding of their evolution and disease ecology. Using a metatranscriptomic approach we identified the viruses in dogs and dingoes from various geographical locations across mainland Australia and sample types, revealing 86 vertebrate-associated viruses belonging to 16 distinct species, including a new vesivirus-like species. Many of the viruses identified here have not previously been sequenced in Australia. We identified important dog pathogens associated with canine infectious respiratory disease syndrome—such as canine pneumovirus, canine herpesvirus, and canine respiratory coronavirus—and gastroenteritis, including canine parvovirus, canine coronavirus, and rotavirus A. The sequences of Australian canine viruses often occupied multiple distinct clades phylogenetically and had little geographic structure, suggesting multiple virus introductions and subsequent spread across the country. Notably, we identified the first RNA virus – rotavirus A – in a dingo. This virus was phylogenetically distinct from dog-associated rotavirus A sequences and more closely related to viruses found in humans and bats, indicative of the past cross-species transmission of a reassortant virus into dingoes, and shows dingoes and domestic dogs may have distinct viromes. Our findings expand the knowledge of viral diversity in Australian canines, improving our understanding of viral movement into and within Australia, as well as the potential zoonotic risks associated with dogs and dingoes.

## Introduction

Australia is host to two taxa of true dogs (tribe Canini)—the domestic dog (*Canis lupus familiaris*) and the dingo (*Canis lupus dingo*). While their taxonomic status and evolutionary history remains contested, it appears that they have arrived separately in Australia. The first species to arrive was the dingo, which is most closely related to New Guinea singing dogs (*Canis lupus hallstromi*) (Fillios and Taçon 2016). The arrival of dingoes likely occurred at least 3,000 years ago, pre-dating the introduction of domestic dog breeds through Western colonisation from the 18th century onwards (Fillios and Taçon 2016). Dingoes can hybridise with domestic dogs, although recent evidence suggests that interbreeding is relatively rare and that behavioural barriers or assortative mating preferences exist (Cairns et al. 2023). Dingoes remain free-living across much of mainland Australia, often coming into contact with humans, wildlife, and domestic dogs (Bombara et al. 2017; Souilmi et al. 2024).

Viruses including canine parvovirus (CPV), canine distemper virus (CDV), and canine respiratory coronavirus (CRCoV) impact dog health in Australia (Kelman et al. 2019; Smith et al. 2021; Wyllie et al. 2016). Given the close proximity dogs share with humans and wildlife, viruses affecting these animals theoretically present a risk of becoming zoonotic or spilling over into wildlife (Ghasemzadeh and Namazi 2015; He et al. 2022). For example, the transmission of CPV between domestic and wild carnivores has been widely reported (Canuti et al. 2022; Kelman et al. 2020b; Nandi and Kumar 2010; Ndiana et al. 2021) and has been associated with clinical symptoms of parvovirus disease (Steinel et al. 2000; Steinel et al. 2001). Wildlife reservoirs of canine viruses are known (Brandell et al. 2021; Canuti et al. 2022) and pose challenges to virus eradication and control measure. Reverse zoonoses events – in which viruses are transmitted from humans to animals – are also of concern, with the transmission of human SARS-CoV-2 to companion animals, including dogs, an important example (Patterson et al. 2020). Moreover, efforts to manage canine viruses, such as those that cause rabies, have substantial economic consequences for pet owners and government agencies (Animal Health Australia 2011). Australia’s aquatic borders and strict biosecurity laws regarding importation present a unique, but currently understudied model, for understanding virus spread and evolution in a relatively isolated locality (Harvey and Foster 2024; Polak 2019).

To date, much of our knowledge of viruses in Australian dogs is restricted to DNA viruses such as CPV (Kelman et al. 2020a). In instances where RNA viruses have been examined, this has been largely limited to clinical and epidemiological investigations without the generation of gene sequence data (Marshall et al. 1984; Wyllie et al. 2016), although a few exceptions exist (Bhatta et al. 2019; Crispe et al. 2011; Moreno et al. 2017; Moreno et al. 2018; Naylor et al. 2001; Naylor et al. 2002; Smith et al. 2021). This knowledge gap is even more apparent in dingoes, with only four cases of CPV reported in dingo puppies and a dingo-dog hybrid (Van Arkel et al. 2019). No metagenomic virus discovery studies of dingoes have been conducted, and it is not currently known if dogs and dingoes share any RNA viruses.

Two metagenomic studies have focused on the faecal virome of Australian shelter dogs with acute diarrhoea (Moreno et al. 2017) and chronic enteropathy (Moreno et al. 2018), identifying numerous DNA and RNA virus families, including the *Astroviridae*, *Coronaviridae* and *Parvoviridae*. However, both studies employed PCR-based sequence-independent, single-primer amplification (SISPA) of viral and genomic DNA. Given the inherent limitations (e.g., uneven and sometimes primer-specific coverage profiles) of this method and the increased potential of artefacts (e.g., repetitive regions) during *de novo* assembly, in most cases viral sequences were not classified at the species level, or made publicly available, nor were phylogenetic analyses conducted (Moreno et al. 2017; Moreno et al. 2018).

Exceptions to this were canine kobu- and astroviruses, where nucleic acids from a single sample identified to have these viruses by SISPA metagenomic sequencing where used to determine the complete genome sequence through Sanger sequencing. These sequences were highly similar to known canine kobu- and astroviruses, sharing 99.41% and 99.14% nucleotide identity with their closest relatives, respectively. In addition, the limited genomic data for many dog viruses globally has hampered our understanding of the patterns of canine viral movement into and within Australia.

Given our limited understanding of canine virus evolution and disease ecology, combined with Australia’s geographic isolation, strict biosecurity measures, and native dingo populations there is clear merit in describing the viruses in domestic dog and dingo populations in Australia. Accordingly, we employed a metatranscriptomic approach (i.e., total RNA-sequencing) to describe the viruses in domestic dog and dingo populations in Australia. We examined multiple tissue types (faecal, tissue, nasal and rectal swabs) across three states—Queensland (QLD), New South Wales (NSW), and Western Australia (WA)— and from different contexts, including dog shelters and rescues, community veterinary clinics, and wild populations. With this data in hand, we investigated virus diversity and evolution within the unique Australian context. In particular, we aimed to: (i) conduct the first metagenomic survey of viruses in dingoes, (ii) compare the viromes of domestic dogs and dingoes to determine the extent of cross-species virus transmission between them, (iii) identify any potential zoonotic pathogens present, and (iv) examine the phylogenetic structure, geographical distribution, and connectivity of Australian canine viruses to infer patterns of virus introduction and spread across the country.

## Methods

### Sample collection and processing

All methods were approved by Taronga Conservation Society Australia’s Animal Ethics Committee (approval number 4c/10/21) and the University of Sydney Animal Ethics Committee (approval number 2022/2096). Carcasses of Myall Lake dingoes were transferred to the Australian Registry of Wildlife Health (the Registry) at Taronga Zoo, Sydney, Australia, for gross and histologic examination, as well as sample collection in accordance with NSW National Parks and Wildlife Act 1974, section 132c, Scientific License number SL100104. Where possible (i.e., not chemically euthanised) carcasses were returned to country for repatriation by Traditional Owners.

### Sample collection and processing

Faeces, rectal and nasal swabs, tissues, and ectoparasites were collected from canines across Australia. Specifically, faecal samples were collected from four dog shelters in NSW: Central West, Illawarra, Coffs Harbour, and Hunter Valley, and one shelter in Perth, WA. Matching sets of nasal and rectal swabs were collected from dogs during three community veterinary clinics in Hope Vale, QLD. Tissue samples were collected from dingoes found deceased or euthanised on welfare grounds from Hawks Nest and Myall Lakes National Park, NSW. Carcasses were stored in a commercial-style chest freezer before being transported to the Registry. The carcasses were allowed to thaw and were necropsied the following day. Tissues were stored at -80 °C. All other sample types were placed in RNAlater. A faecal sample was also opportunistically collected from Myall Lakes National Park (-32.626E, 152.201S). Trail cameras (browning strike force HD) were deployed at dingo scent-marking sites in Myall Lakes National Park as part of a separate study. This faecal sample was collected at one of these sites 21 hours after a dingo was seen on a trail camera displaying behaviours consistent with having just scent marked (slightly off camera), in the exact location the sample was collected and stored in a portable -80°C freezer. Additional dingo faecal samples were collected from a dingo rescue centre in WA and transported on ice to laboratory facilities. All samples were then stored at -80°C until RNA extraction. Dog age, sex, and general health status information were recorded by technicians for the shelter samples and by a veterinarian for the Hope Vale samples. Dog ages were classified into three categories: puppies (less than 1 year old), young adults (1–2 years old), and adults (over 2 years old). Most animals showed no obvious signs of pathology. Those with signs of pathology are recorded in Supplementary Table 1.

### RNA extraction and metatranscriptomic sequencing

Individual aliquots of each sample were placed into 600 μl of lysis buffer containing 1% β-mercaptoethanol (Sigma-Aldrich). Faecal samples and swabs were homogenised in a QIAshredder column (Qiagen). Dingo tissue was homogenised using a TissueRuptor (Qiagen) at a speed of 5,000 rpm for up to one minute with 0.5% antifoaming reagent (Reagent DX, Qiagen) added to the lysis buffer. RNA was then extracted from the supernatant using the RNeasy Plus Mini Kit (Qiagen), following the manufacturer’s protocol. Extracted RNA was pooled in roughly equimolar ratios, grouping by location, sample type, sex, condition, and age. This resulted in 70 pools with a median of 2.5 samples per pool (minimum = 1, maximum = 6) (Supplementary Table 1). Sequencing libraries were constructed using the Truseq Total RNA Library Preparation Protocol (Illumina). Host ribosomal RNA (rRNA) was depleted using the Ribo-Zero Plus Kit (Illumina) and paired-end sequencing (150 bp) was performed on the NovaSeq 6000 platform (Illumina). Library construction and sequencing were performed by the Australian Genome Research Facility (AGRF) in Melbourne, Australia. Seventeen of the Hope Vale libraries were sequenced over two lanes. Each library generated from a lane was processed through the pipeline and analysed individually. When assembling final genomes and estimating abundance reads from both libraries, they were used as a single input.

### Identification of novel virus sequences

Sequencing libraries were processed following the BatchArtemisSRAMiner pipeline (v1.0.4) (Mifsud 2023). Reads underwent quality trimming and adapter removal using Trimmomatic (v0.38) with parameters SLIDINGWINDOW:4:5, LEADING:5, TRAILING:5, and MINLEN:25, prior to assembly (Bolger et al. 2014). *De novo* assembly was conducted using MEGAHIT (v1.2.9) (Li et al. 2015). Assembled contigs were compared to the RdRp-scan (RNA-dependent RNA polymerase) RdRp core protein sequence database (v0.90) (Charon et al. 2022) and the protein version of the Reference Viral Databases (v26.0) (Bigot et al. 2019; Goodacre et al. 2018) using Diamond BLASTx (v2.1.6) (Buchfink et al. 2021) with an e-value cut-off of 1 × 10^−4^ for the RdRp-scan database and 1 × 10^−10^ for the RVDB database. To exclude potential false positives, contigs with hits to virus sequences were used as a query against the NCBI nucleotide database (as of December, 2023) using BLASTN (Camacho et al. 2009) and the NCBI non-redundant protein (nr) database (as of December, 2023) using Diamond BLASTx. We used sequence similarity to known vertebrate viruses (e.g., % amino acid similarity), and in some cases phylogenetic grouping, to determine if a virus was potentially vertebrate infecting. Viruses identified as likely of non-vertebrate origin were excluded from further analysis.

To remove viruses present as a result of index-hopping from another library, we excluded those that meet the following three criteria: (i) sequenced on the same lane, (ii) the total read count was <0.1% of the read count in the other library, and (iii) were >99% identical at the nucleotide level. To screen for endogenous viral element (EVEs) within the viral-like contigs, all putative virus-like nucleotide sequences were compared to the corresponding host genome (*Canis lupus familiaris* or *Canis lupus dingo*) using the BLASTN algorithm with an E-value cutoff of 1 × 10^−20^. In addition, all virus-like sequences were checked for host gene contamination using the contamination function implemented in CheckV (v0.8.1) (Nayfach et al. 2021). No EVEs were detected.

To infer whether a sequence represents a novel virus we used the demarcations set by the International Committee of Viral Taxonomy (ICTV) for each virus family. Where no demarcations were provided, we classified a virus as novel if it shared <95% amino acid identity across the RdRp and represented a distinct clade.

### Genome extension, annotation and abundance

To potentially extend contigs and check for areas of heterogeneous coverage, sequence reads were mapped onto virus-like contigs with BBMap (v37.98) (Bushnell 2014). For known dog viruses, the closest full-genome representative sequence as determined by web BLASTN was used as a reference for mapping. The resulting library consensus was then remapped with 98% nucleotide identity. ORFs were predicted using the Geneious Prime Find ORFs tool (v2022.0) (https://www.geneious.com/ ) (Kearse et al. 2012). Protein functional domains were annotated using the InterProScan software package (v5.56-89.0) (Jones et al. 2014) with the TIGRFAMs (v15.0), SFLD (v4.0), PANTHER (v15.0), SuperFamily (v1.75), PROSITE (v2022_01), CDD (v3.18), Pfam (v34.0), SMART (v7.1), PRINTS (v42.0), and CATH-Gene3D databases (v4.3.0). Classification of Rotavirus A (RVA) genomes was performed using the Rotavirus A Genotyping Tool Version 0.1 (https://www.rivm.nl/mpf/typingtool/rotavirusa/) (check Supplementary Table 2 for detailed results).

The abundance of each virus contig was calculated using Bowtie2 (v2.2.5) (Langmead and Salzberg 2012). Trimmed reads were mapped against each virus contig at 99% identity. Samtools (v1.6) (Li et al. 2009) was used to calculate read coverage. Abundance was reported as a percentage of the total number of reads mapped divided by the total number of trimmed reads in that library.

### Assessment of sequencing library composition

Library composition was assessed by Kraken2 (v2.1.2) (Breitwieser et al. 2018; Wood et al. 2019) for the purpose of confirming host species and investigating pathogen co-occurrence in libraries in which we identified canine infectious respiratory disease syndrome associated viruses. Kraken2 was ran on the trimmed reads using non-standard options --minimum-hit-groups 3 and --report-minimizer-data against the nt Kraken data set 5/30/2024 obtained from https://benlangmead.github.io/aws-indexes/k2. Bracken (v2.9) (Lu et al. 2017) was then used to estimate the abundance of each species and KrackenTools (v1.2) (Lu et al. 2022) and Krona (v2.8.1) (Ondov et al. 2011) were used to process and visualise the output.

### Phylogenetic analysis

To determine the evolutionary relationships of the viral sequences obtained to known viruses, we inferred maximum likelihood phylogenetic trees derived from nucleotide and amino acid multiple sequence alignments. The sequence type (nucleotide or amino acid) and region (RdRp or capsid) used for the alignment were dependent on standard practice for the virus group in question, such as species demarcation criteria established by the ICTV and the degree of sequence similarity between the identified and reference sequences. For example, when analysing the phylogenetic relationships within a single virus species (e.g., canine associated *Sapporo virus* sequences), nucleotide sequences from the capsid region were typically used, as the amino acid sequences of RdRp tend to be nearly identical. Sequences determined in this study were aligned with reference sequences using MAFFT (v7.402) (Katoh and Standley 2013) using the E-INS-I algorithm. Ambiguously aligned regions were removed using trimAl (v1.4) (Capella-Gutiérrez et al. 2009), employing variable conservation thresholds (i.e., minimum percentage of alignment columns to retain) and gap thresholds (i.e., the minimum fraction of sequences without a gap needed to keep a column), as well as the automated parameter selection method gappyout.

Phylogenetic trees were estimated using the maximum likelihood method in IQ-TREE 2 (v2.1.7) (Minh et al. 2020) with the selection of the best-fit substitution model determined using ModelFinder (Kalyaanamoorthy et al. 2017). The sequence type, trimAl parameters and IQ-TREE 2 substitution model selected are presented in Supplementary Table 3. Branch support was calculated using 1,000 bootstrap replicates with the UFBoot2 algorithm and an implementation of the SH-like approximate likelihood ratio test within IQ-TREE 2. Phylogenetic trees were annotated using the R packages ape (5.7.1) (Paradis and Schliep 2019). Phytools (2.1.1) (Revell 2024) and ggtree (v3.8) (Yu et al. 2017), and further edited in Adobe Illustrator.

### Statistical comparisons of virus count, diversity, and abundance

To examine how virus count, diversity, and abundance varied across sample type, host sex, and host age, we conducted Kruskal-Wallis tests using the stats package (v4.2.3) in R. Virus count was defined as the number of dog-associated viruses per library, while diversity was virus count summarised at a species level as to count only one virus per species. Abundance was measured as the percentage of total library reads assigned to that virus. For visualisation, boxplots were generated using the ggplot2 package (v3.5.1) in R.

### Statistical comparisons of genetic and geographical distances

To determine the relationship between virus sequence relatedness and geographical distance, we performed Mantel tests for each virus species. As they are so closely related, canine vesivirus (CaVV) and the canine vesi-like viruses identified in our study were treated as a single species. Pairwise sequence identity percentages across the genome were obtained using BLASTN. In the case of segmented viruses only the RdRp-containing segment was used. Genetic distance was calculated by subtracting the pairwise sequence identity (%) from 100. Geographical distances between sampling locations were calculated using the Haversine formula using the distHaversine function from the geosphere package (v1.5-18) in R. Mantel tests were conducted using Spearman’s rank correlation coefficient with 999 permutations through the mantel function from the vegan package (v2.6-4) in R.

## Results

We characterised the virome of domestic dogs and dingoes sampled from locations across Australia. Total RNA-sequencing was performed on 185 samples pooled into 70 libraries (87 when considering the Hope Vale libraries sequenced on two lanes), grouping samples by location, sample type, sex, condition, and age (Figure 1). A total of 8,292,278,589 sequence reads were generated with a median of 155,755,544 (range 63,732,230 – 237,451,130). *De novo* assembly of the sequencing reads resulted in a median of 144,407 (range 251 – 738,858) contigs with a range of per library, for a total of 19,399,943 contigs. Library statistics are provided in Supplementary Table 4.

**Figure 1.**
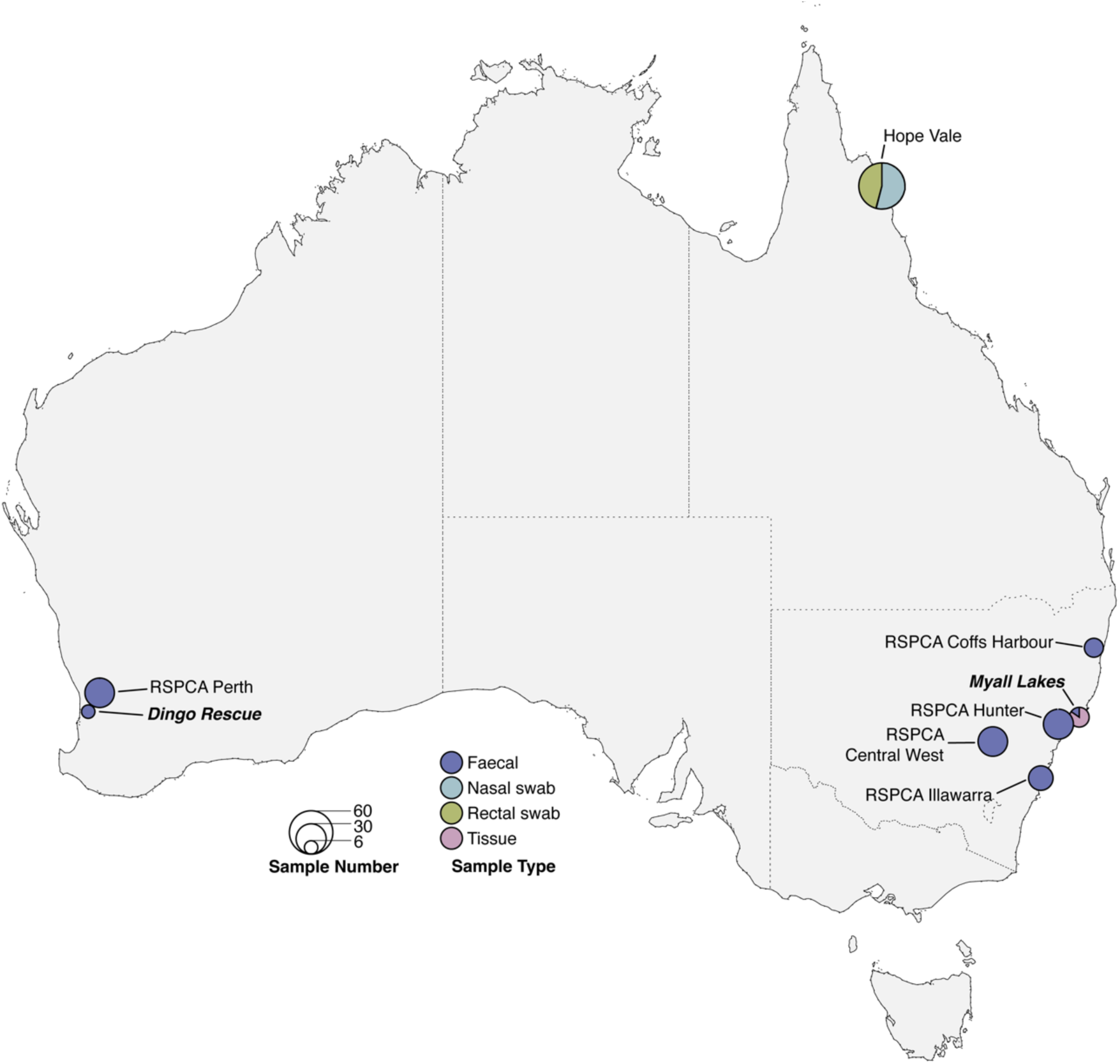
Map showing the composition and number of the dog and dingo samples across mainland Australia. The pie chart reflects sample type, and the size reflects the number of individual samples collected at each location. Sample sites where dingoes were collected are labelled in italics.

### Identification of potential dog and human pathogens in Australia

We focused our analysis on vertebrate infecting viruses, excluding those that are likely associated with diet or gut flora such as picobirnaviruses that are likely associated with bacteria (Sadiq et al. 2024). Overall, we detected 86 vertebrate-associated viruses, 82 of which were RNA viruses and 4 DNA viruses. These viruses were classified into 16 virus species, 15 of which are known dog viruses as well as one novel species in the *Caliciviridae* family of RNA viruses. Members of the RNA virus order *Picornavirales* were the most frequent both in terms of the number of distinct species (n = 6) and the number of libraries (n = 34) (Supplementary Table 5). Members of the family *Picornaviridae* comprised a range of genera: *Kobuvirus*, *Dicipivirus*, *Mischivirus*, and *Sapovirus*. The remaining viruses were members of the *Caliciviridae, Coronaviridae*, *Pneumoviridae*, *Astroviridae*, and DNA viruses from the *Parvoviridae* and *Orthoherpesviridae*. In contrast to many viral metagenomic studies in other mammals, nearly all the viruses discovered here are known rather than novel.

The number of canine viruses, their diversity in terms of the number of species, and their abundance varied across sample type. In particular, faeces contained significantly higher canine virus numbers (H (2) = 17, P = 0.0002), diversity (H (2) = 19.7, P = 0.00005) and abundance (H (2) = 19.7, P = 0.00001) than other sample types. There was also a significant difference in canine virus abundance between sexes (H (1) = 5.353, P = 0.02) with males having a higher overall abundance. No significant difference was observed in the number or diversity of canine viruses. There was no detectable relationship between these variables and dog age (Figure 2, Supplementary Table 6).

**Figure 2.**
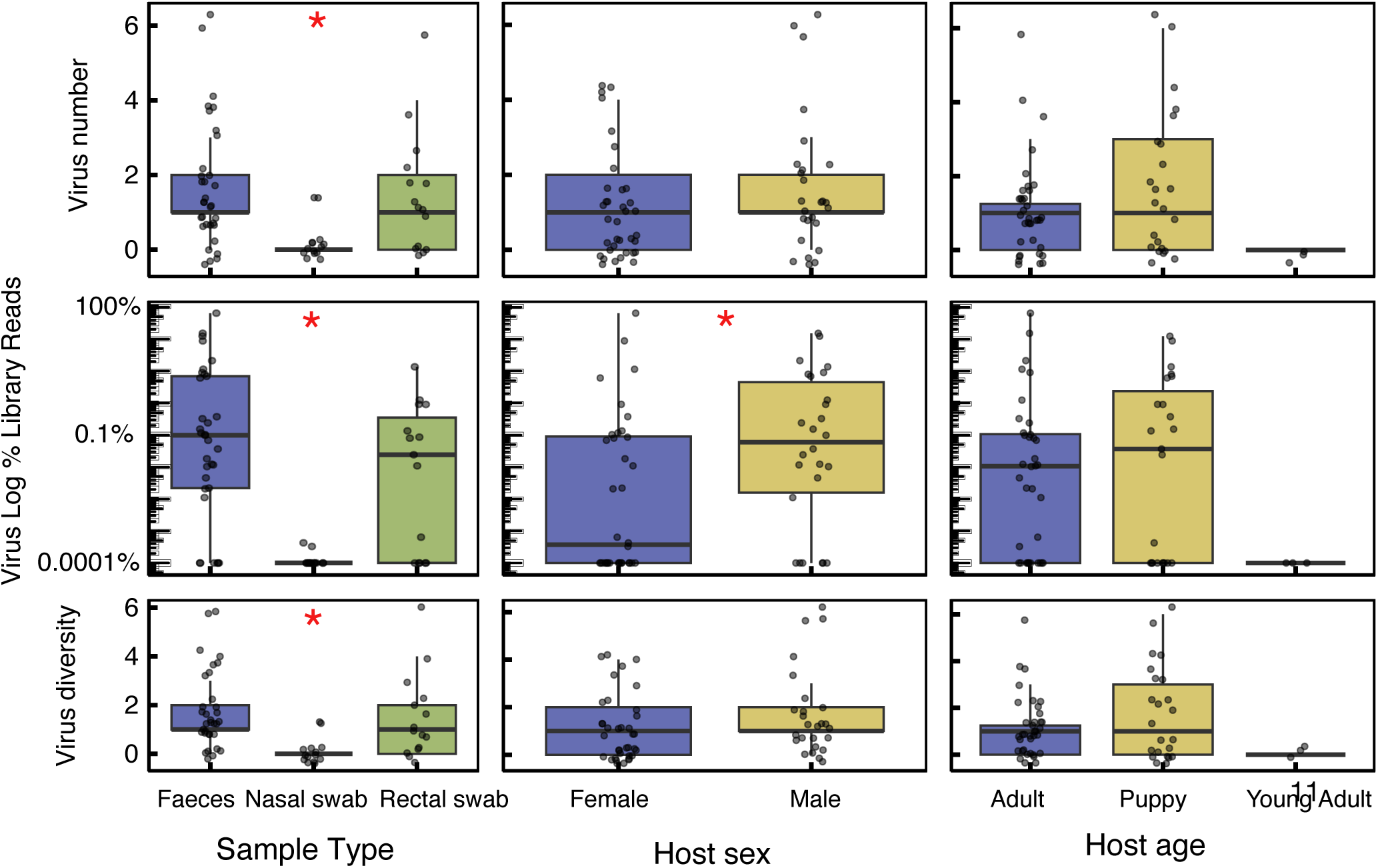
Boxplot showing the differences in virus number, percentage of total library reads and virus diversity between sample type (faeces, nasal swab, and rectal swab), host sex (female and male) and host age (adult, young adult, and puppy). Significant differences between the groups as determined by a Kruskal-Wallis rank sum test are shown with a red asterisk. In the boxplots, the middle lines show the median, and upper and lower hinges show the first and third quartiles.

We now describe specific virus groups in turn.

#### Canine infectious respiratory disease syndrome (CIRDS)

This is a syndrome associated with respiratory symptoms and collectively described as “kennel cough”. It is associated with multiple bacterial and viral pathogens including *Bordetella bronchiseptica*, CRCoV, Canine pneumovirus (CnPnV), CDV, and Canine herpesvirus (CHV, canid alphaherpesvirus 1). We identified fragments of CnPnV, CRCoV and CHV in three sequencing libraries (Supplementary 4). A 369 bp fragment of CnPnV was assembled from hopevale_40, a library comprising a single nasal swab from a female puppy, at a relatively low abundance of 0.36 fragments per kilobase of transcript per million mapped reads (FPKM) and representing 4.27 x 10^-6^ % of total library reads. The associated veterinary report described the dog as “mangey, thin, kennel cough”. This fragment was most closely related to a virus associated with a 2012 CIRDC outbreak in Italy (NC_025344, 98.9% nucleotide identity) (Decaro et al. 2014). Phylogenetic trees inferred from the partial RdRp fragment placed this virus with the European Bari/100-12 sequence rather than those identified from outbreaks in Thailand or the US (Piewbang and Techangamsuwan 2019; Thieulent et al. 2024) (Figure 3a).

**Figure 3.**
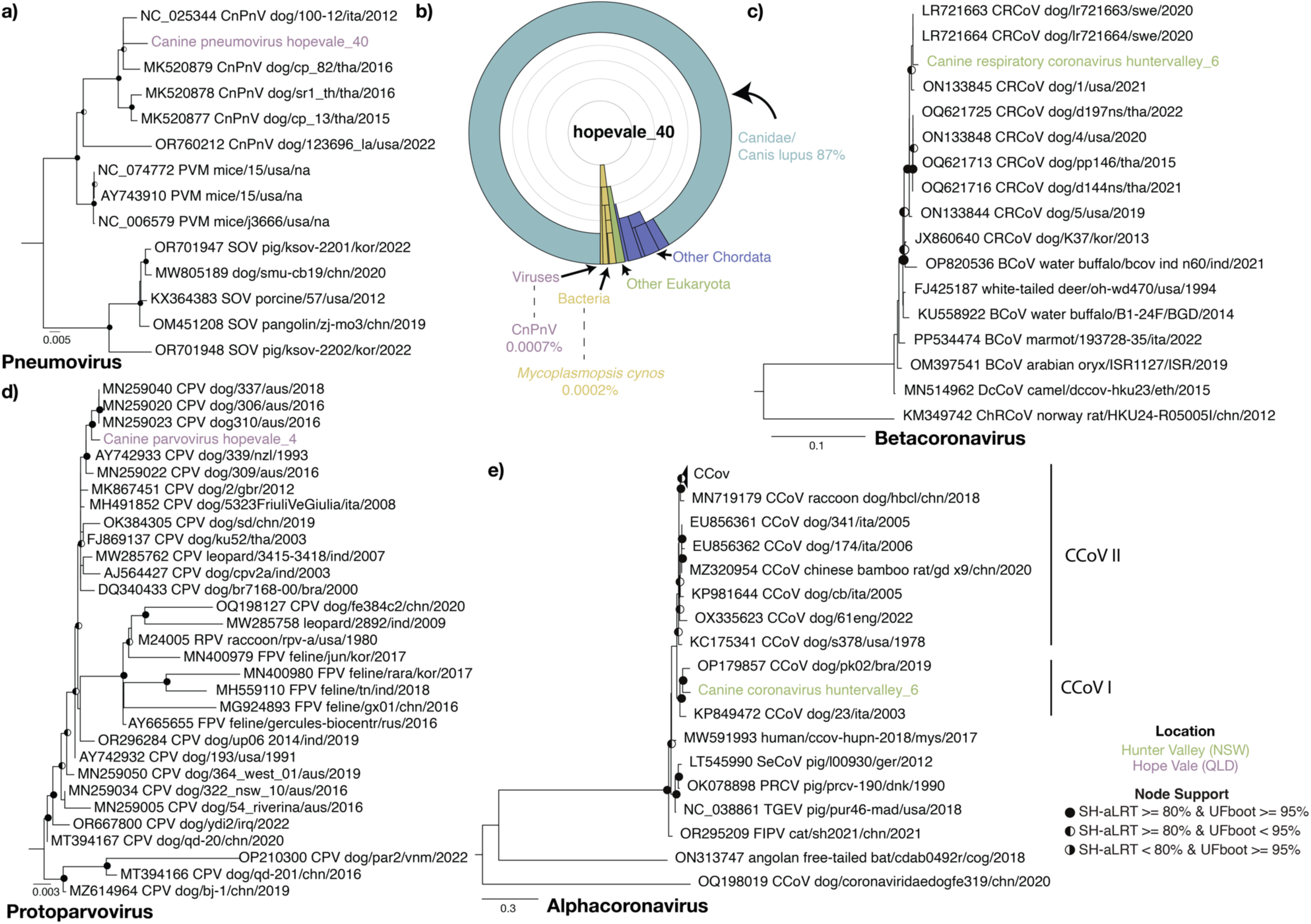
(a,c-e) Maximum likelihood phylogenetic trees of four viral pathogens. Phylogenies were inferred using nucleotide sequences from the (a) *Pneumovirus* RdRp gene, (c) *Betacoronavirus* RdRp, (d) *Protoparvovirus* VP1, and (e) *Alphacoronavirus* ORF1b. The trees are midpoint rooted for clarity with the exception of the protoparvoviruses which are rooted on the node leading to OP210300, MT394166, MZ614964. Branch lengths reflect the number of substitutions per site. Strong node support (SH-aLRT >= 80% and UFboot >= 95%) is indicated by a black circle while intermediate support (SH-aLRT <80% and UFboot >95% or SH-aLRT >80% and UFboot <95%) is indicated by a half-filled circle. The tip labels of the sequences identified in this study are coloured by sampling location. (b). Krona chart showing the classification of reads associated with dogs and other eukaryotes as well as the potential agents of canine infectious respiratory disease syndrome.

To examine whether any other pathogens were present in the hopevale_40 library, rRNA-depleted reads were assessed for taxonomic associations using Kraken. This revealed that most of the bacterial reads identified were associated with Mycoplasmatota (83% of bacteria, 2% of library), and a small percentage (0.01%) of these were associated with *Mycoplasmopsis cynos,* part of the CIRDS complex. The library abundance of *Mycoplasmopsis cynos* was slightly less (0.0001%) than that of CnPnV (0.0007%). However, the majority of the Mycoplasmatota reads assigned at the species level were from *Mycoplasma cavipharyngis* (38%) and *Mycoplasmoides fastidiosum* (12%), which are not associated with disease (Figure 3b). No other dog pathogens were detected, and no CnPnV was found in the associated rectal swab from this individual.

Fragments of CRCoV, a *Betacoronavirus* associated with CIRDS, were assembled from the huntervalley_6 library that comprised five adult male dog faecal samples. There were no reports of dogs with respiratory symptoms or general illness. Both fragments were at low abundance, representing 0.24 and 0.32 FPKM and 3.14 x 10^-6^ and 5.23 x 10^-6^ percentage reads, respectively. Interestingly, there was only a single nucleotide change between these two fragments, 339 and 388 bp in length, and a sequence (LR721664.1) associated with a *Betacoronavirus* associated CIRDS outbreak in Swedish dogs, that was thought to have entered Sweden in approximately 2010 and has since become endemic (Wille et al. 2020). Phylogenies inferred from the RdRp region support this close association with the Swedish sequences (Figure 3c).

Fragments of CHV, an alphaherpesvirus, were identified in the hopevale_21 library which comprised nasal swabs from three adult female dogs without respiratory symptoms. The fragments shared 99-100% nucleotide identity with CHV sequence V1154 (KT819631.1). Phylogenetic analysis of the polymerase fragment placed this sequence within the larger diversity of CHV, although there was insufficient sequence to reliably infer evolutionary history and whether it was related to other Australian sequences.

#### Gastrointestinal viruses

Gastroenteritis in dogs has been linked to a number of viruses including CPV, Canine coronavirus (CCoV), Rotavirus A (RVA), and potentially Canine vesivirus (CaVV) and Canine astrovirus (CaAstV) (Martella et al. 2012; Renshaw et al. 2018). RVA is a leading cause of acute diarrhoea in humans, and dog-associated sequences have been associated with zoonotic transmission to humans. Notably, we identified three species of these known pathogens in seemingly healthy dogs.

The complete genome of CPV was assembled in the hopevale_4 library that comprised rectal swabs from three male puppies with no signs of disease. The sequence represents 0.003% of total library reads. Across the VP1 and NS1 genes this sequence shared the highest nucleotide identity with those from New Zealand in 1993 (AY742933). Indeed, phylogenetic analysis of the VP1 placed this sequence with sequences sampled in 2016 and 2018 from NSW in what is termed as the CPV-2a clade 5 (Kwan et al. 2021) (Figure 3d).

CCoV was assembled from the huntervalley_6 library which comprised a pool of 5 adult male dog faecal samples who were seemingly healthy, and represented 0.34% of total library read abundance. This sequence is distinct from the previously identified CCoVs in Australia, sharing 70%, 60%, and 50% amino acid identity in the spike to the QLD CCoV-IIb sequence (MW383487) (Smith et al. 2021) and two divergent NSW sequences [AF516907 (Naylor et al. 2002) and AF327928 (Naylor et al. 2001)], respectively. Phylogenetic analysis of huntervalley_6 ORF1b placed huntervalley_6 in CCoV I (Figure 3e). As with other CCoV I sequences, an additional accessory protein gene ORF3 was predicted between nucleotides 25,518 and 26,141.

Coding complete genomes of RVA were recovered in four libraries from the Hunter Valley (n = 4) and Central West shelter (n = 1), in both puppies and adults. Of particular note, partial VP1 (407 bp), NSP1 (317 bp) and NSP3 (280 bp) sequences were also recovered in the library of a dingo from Myall Lakes National Park that comprised a single faecal sample.

With the exception of the fragments in the myalllakesdingo_4 library (7.55 x 10^-6^%), these viruses were very highly abundant, on average representing 22% of library reads (min = 0.015%, max = 64.45%). Viral abundance was also higher in adult dogs than puppies. Despite this, both the dogs and dingoes were reported to be healthy.

The RVA sequences were classified using an approach used by the Rotavirus Classification Working Group (RCWG) based on the 11 genome segments (Matthijnssens et al. 2011). Accordingly, our sequences were classified as ‘G3-P[3]-I3-R3-C3-M3-A9-N2-T3-E3-H6’. The Myall Lake dingo sequences were also classified as R3 (VP1) and T3 (NSP3), but typing could not be performed with the NSP1 fragment. This classification was supported by phylogenetic analysis of the VP4 gene, which placed the sequences in a monophyletic group with other members of the same genotype identified in dogs from Thailand (Figure 4a). As no VP4 or VP7 sequences were recovered for the myalllakesdingo_4 RVA, a phylogenetic analysis was performed using VP1 (Figure 4b). This revealed that the Myall Lakes dingo RVA was distinct from the those identified in dogs, sharing ∼90% nucleotide identity across the fragments (Figure 4a). Instead, notably, the dingo RVA VP1 sequence grouped with a human associated RVA sequence (KU597747) identified from a child with diarrhoea from China in 2014 which is thought to have originated from a reassortment from bat MYAS33 or simian rotaviruses (Dong et al. 2016). This similarity to non-canine associated sequences was also apparent in the other two fragments recovered in this library, with all three fragments having closest nucleotide matches to *Myotis horsfieldii* (OR868019, OR868019) and *Rhinolophus affinis* (OR868034) RVA sequences, identified in bats from China in 2016 and 2017, respectively.

**Figure 4.**
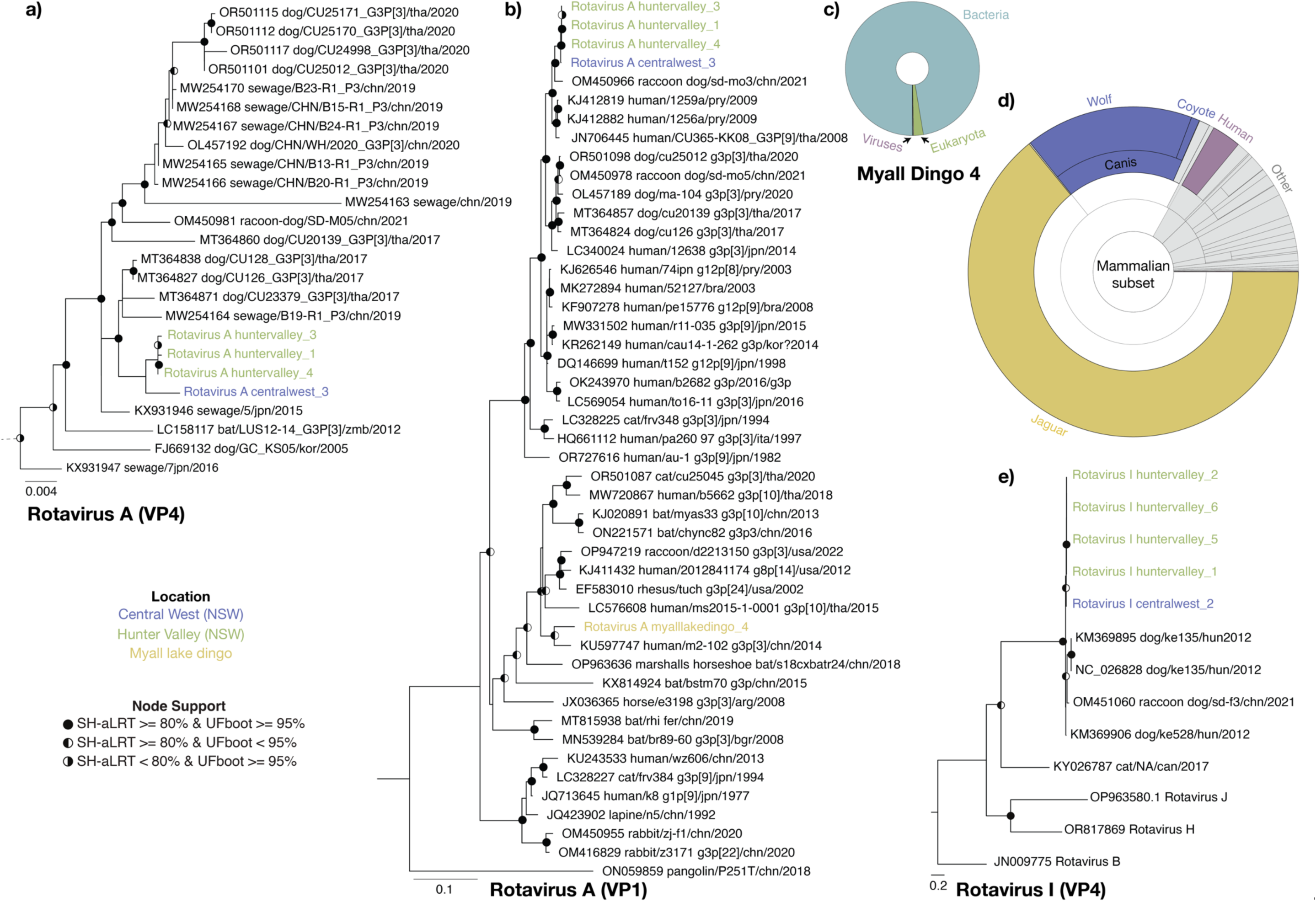
Maximum likelihood phylogenetic trees of canine rotaviruses. Phylogenies were inferred using (a) Rotavirus A (RVA) VP1, (b) RVA VP4, and (c) Rotavirus I VP4. The trees are midpoint rooted for clarity, with branch lengths reflecting the number of substitutions per site. Strong node support (SH-aLRT >= 80% and UFboot >= 95%) is indicated by a black circle, while intermediate support (SH-aLRT <80% and UFboot >95% or SH-aLRT >80% and UFboot <95%) is indicated by a half-filled circle. (b) RVA VP4 is a clade extracted from a larger G3P3 RVA phylogeny available in the GitHub repository (https://github.com/JonathonMifsud/Australian-Canine-Virome/). The tip labels of sequences identified in this study are coloured by sampling location. (d) Krona chart showing the classification of reads associated in the myalllakedingo_4 library, Super kingdom level classification of Bacteria, Viruses and Eukaryota are labelled. (e) mammalian subset of the myalllakedingo_4 library composition as shown in d.

The taxonomic composition of the myalllakedingo_4 library was further analysed to check for evidence of another possible source of RVA origin (dietary or contamination). The myalllakedingo_4 library was predominately comprised of bacteria (97%). Of the eukaryotic reads detected in this library, 57% were associated with Arthropoda, 21% of which were associated with the dung flies (*Sepsis cynipsea*) (Figure 4c). Mammalian reads, which represent just 0.3% of all reads, were 64% *Panthera onca* (jaguar) and 16% *Canis lupus* (Figure 4d). To examine this further, we used BLASTN to compare the myalllakedingo_4 contigs to the nucleotide database and found that 74% of the vertebrate associated contigs with an abundance of >= 50 reads were associated with *Canis*. Notably, contigs with top hits to *Panthera onca* were associated with the reference *Panthera onca* voucher GTJAG006 16S ribosomal RNA gene (KX419654) which, when further examined, was found to be a misannotated *Fusobacteriaceae* sequence. No human associated contigs were found to have >40 reads. A similar library composition was observed in the myalllakedingo_5 library, with 92% of sequence reads assigned to bacteria and 0.6% assigned to the genus *Canis*. In this case, the faecal sample was collected directly from the individual during dissection under sterile conditions.

A rarely detected rotavirus species, Rotavirus I (RVI), was recovered from five libraries from the Hunter Valley (n = 4) and Central West shelter (n = 1). Fragments were also detected in centralwest_4 library, but no VP1 or VP4 sequences were recovered. There was no clear link between the presence of this virus and pathology in the dogs (Supplementary Table 1). RVI sequences formed a single clade as a sister group to the known RVI sequences (Figure 4e), and there was very little diversity between our sequences (99.4% average nucleotide identity based on a multiple sequence alignment of the four sequences). RVI abundance varied between 0.0001% to 0.77% of total library reads, with an average of 0.19%.

Sequences related to CaVV were detected in faeces and rectal swab samples from 12 libraries. There was no association between the presence of CaVV and pathology in the dogs (Supplementary Table 1). Phylogenetic analysis of the VP1 demonstrates that Australian CaVV sequences formed multiple independent lineages within the diversity of this virus, including a distinct clade that was associated with Illawarra and Central West shelter dogs (Figure 5b). This clade appears divergent from known CaVV sequences, sharing ∼72% amino acid identity with its closest relative from a canine nasal swab taken in China in 2024 (XHB18322). Consequently, this clade likely represents a novel species within the genus *Vesivirus*.

**Figure 5.**
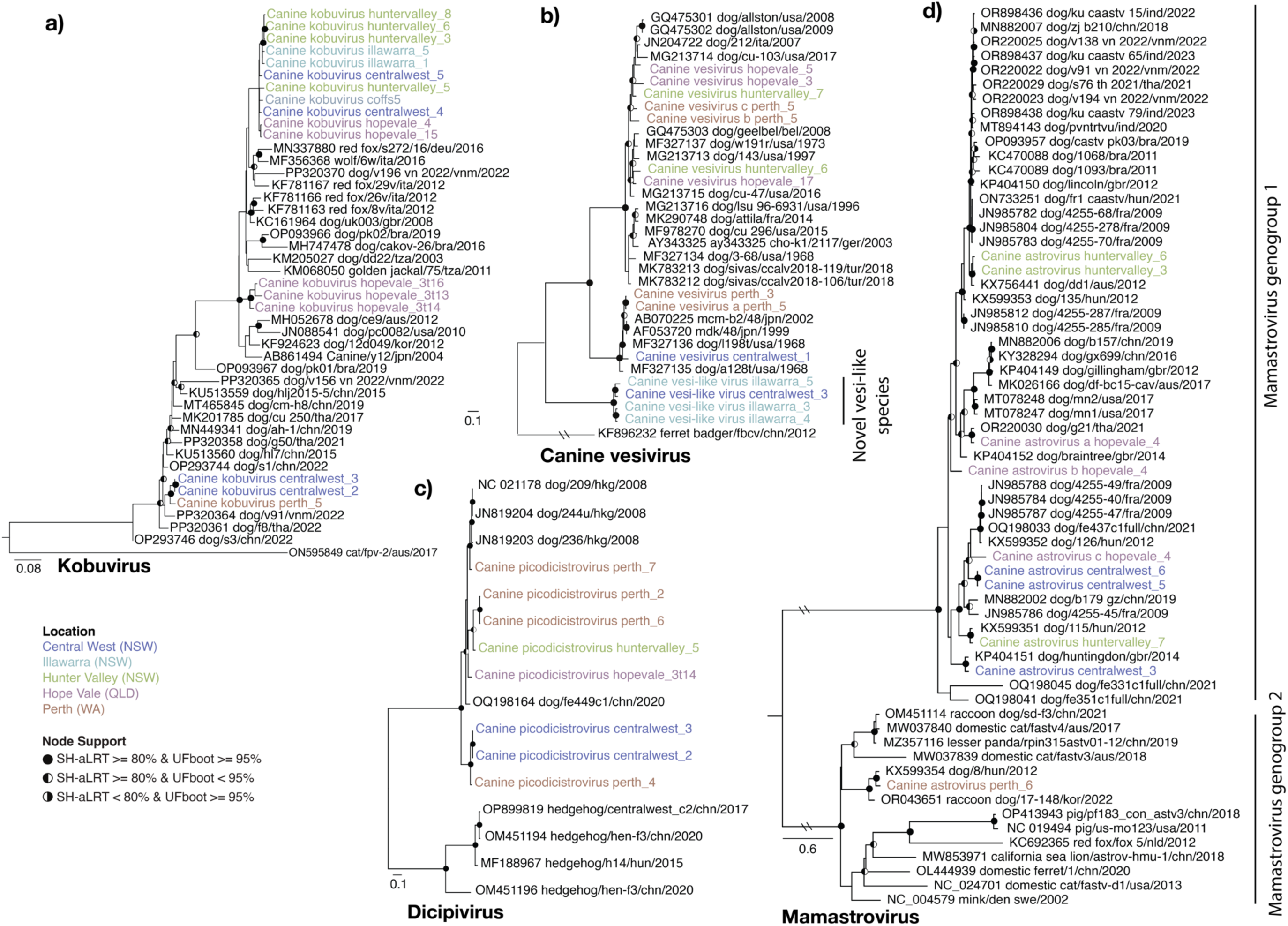
Maximum likelihood phylogenies of kobuviruses, vesivirus, dicipivirus and mamastroviruses. Phylogenies were inferred using (a) Kobuvirus VP1 nucleotide, (b) Canine vesivirus VP1 amino acid, (c) Dicipivirus VP1 amino acid, and (d) Mamastrovirus capsid nucleotide sequences. The trees are midpoint rooted with the exception of Canine vesivirus which is rooted on the Ferret badger calicivirus (KF896232) outgroup, the sequence of which has for been removed for visual clarity. Branch lengths reflect the number of substitutions per site. Strong node support (SH-aLRT >= 80% and UFboot >= 95%) is indicated by a black circle while intermediate support (SH-aLRT <80% and UFboot >95% or SH-aLRT >80% and UFboot <95%) is indicated by a half-filled circle. The tip labels of sequences identified in this study are coloured by sampling location. The two lines on the branches of the Mamastrovirus tree indicate that these branches were shortened for visualisation purposes.

### Co-occurrence of viruses in faecal libraries

Cases of gastroenteritis are commonly characterised by coinfections between multiple viruses, with seemingly weak or unknown associations with disease. In addition to the viruses described above, we detected the following viruses in faeces or rectal swab libraries: Canine kobuvirus (CaKoV) (n = 17 libraries), CaVV/CaVV-like (n = 12), Canine picodicistrovirus (CPDV) (n = 10), CaAstV (n = 8), sapovirus (SaV) GXIII (n = 7), Norovirus (NoV) (n = 6), Canine picornavirus (CanPV) (n = 5) and ucalufa virus (n = 1) (Figure 5, 6). In these libraries the presence of a second dog virus was detected ∼50% of the time (19/38 libraries) and the average number of co-occurring viruses per library was 2.02 (range 1 - 6). Six viruses were found in HU6 and CW3, the former containing CaAstV, CaCV, CCoV, CRCoV, and RVI. There was no relationship between the number of co-occurring viruses and the number of individual samples per library nor sampling location. CaKoV frequently co-occurred with other dog viruses (14/17) and was rarely the only virus found in a library. Notably, where noroviruses and SaVs were identified, CaKoV was also detected 83% and 71% of the time (Figure 7).

**Figure 6.**
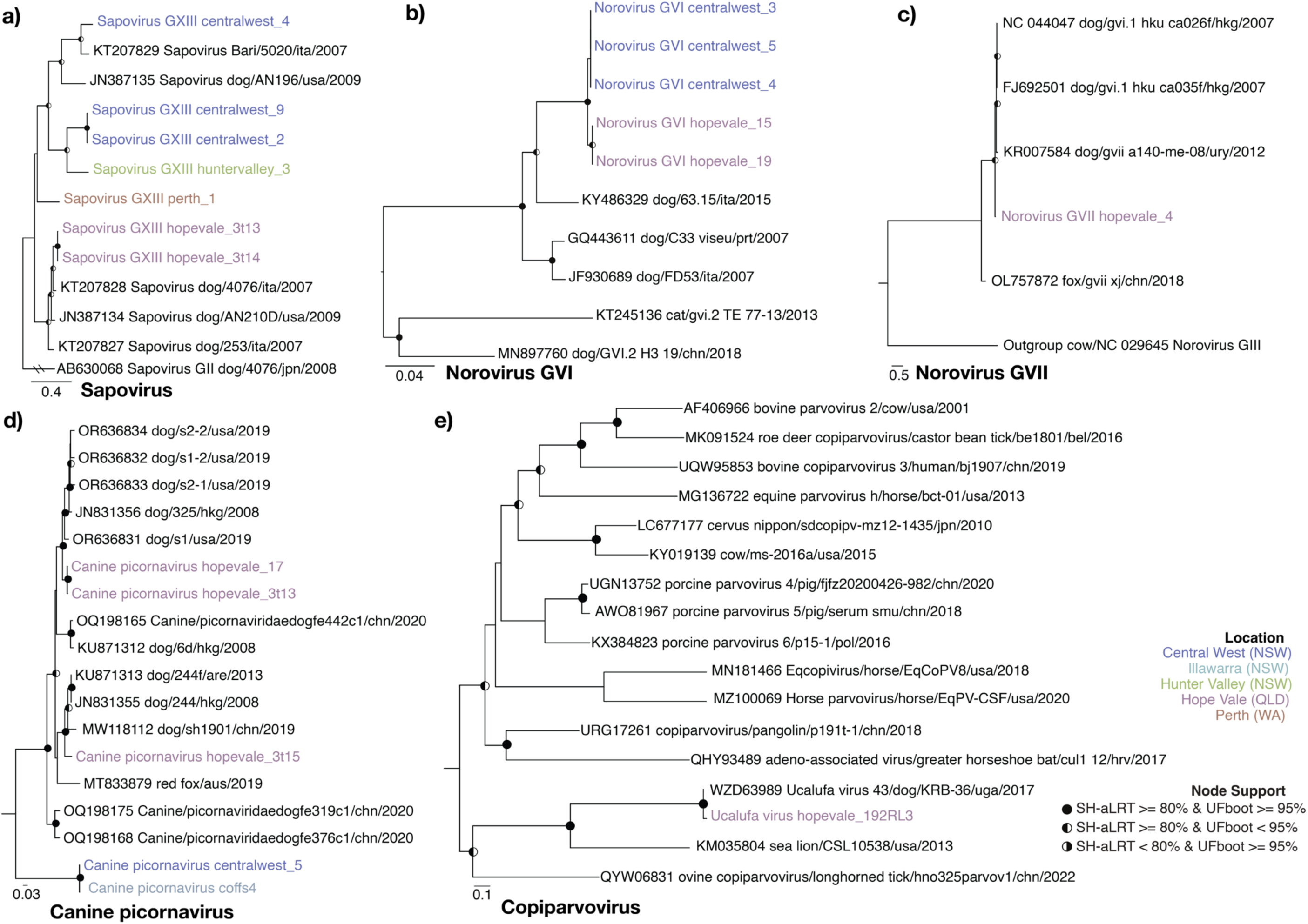
Maximum likelihood phylogenies of sapoviruses, noroviruses, picornaviruses and copiparvoviruses. Phylogenies were inferred using (a) Sapovirus VP1 nucleotide, (b) Norovirus GVI VP1 nucleotide, (c) Norovirus GVII VP1 nucleotide, (d) Canine picornavirus RdRp nucleotide, and (e) Copiparvovirus RdRp amino acid. The trees are midpoint rooted for clarity, with branch lengths reflecting the number of substitutions per site. Strong node support (SH-aLRT >= 80% and UFboot >= 95%) is indicated by a black circle while intermediate support (SH-aLRT <80% and UFboot >= 95% or SH-aLRT >80% and UFboot <95%) is indicated by a half-filled circle. The tip labels of sequences identified in this study are coloured by sampling location. The two lines on the branches of the Sapovirus tree indicate that these branches were shortened for visualisation purposes.

**Figure 7.**
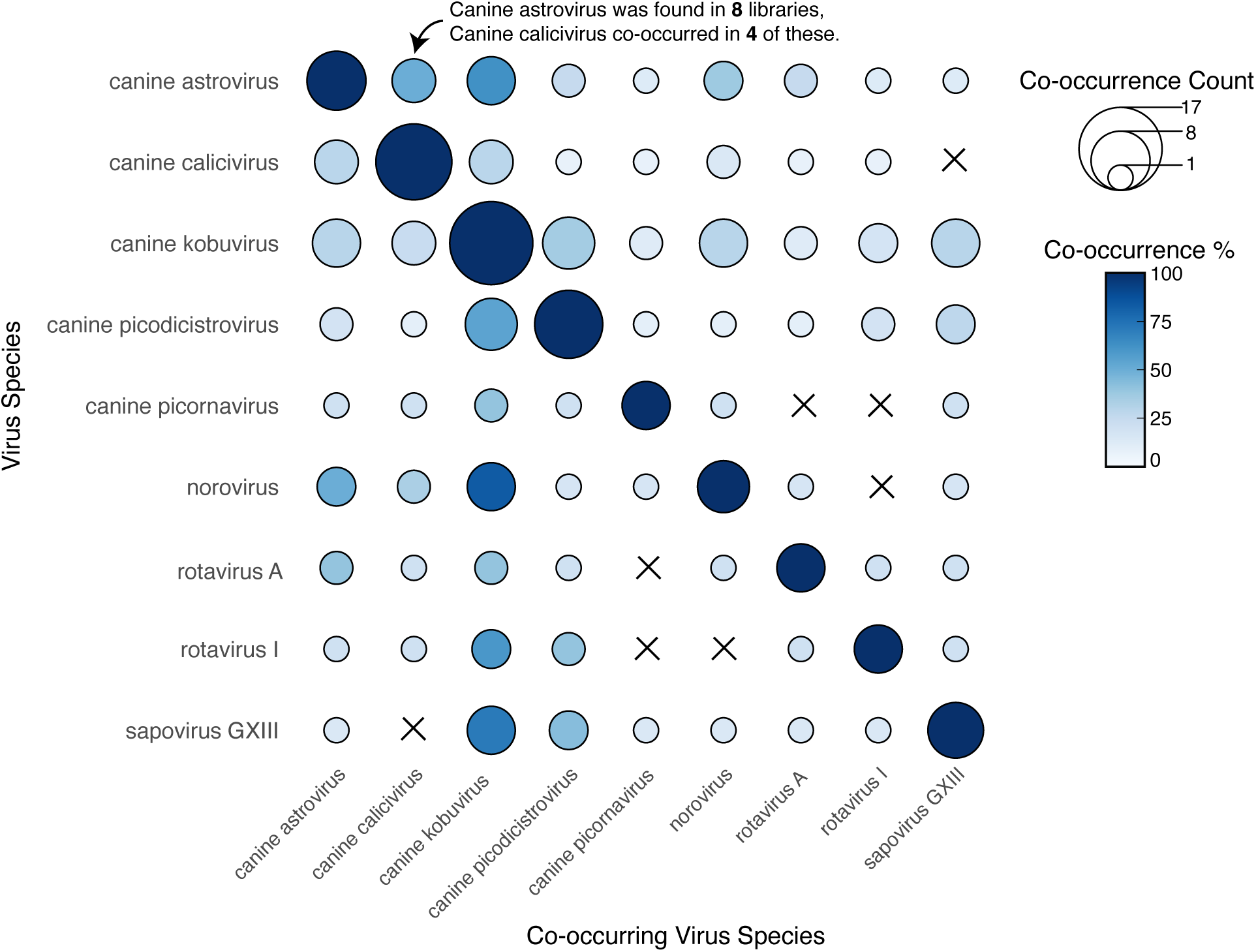
Virus co-occurrence heatmap matrix. For virus species that occur more than once, the frequency and percentage at which they are found with other viruses within a given library is shown. The size of the circle indicates the number of occasions while the colour represents the percentage of time the “Co-occurring Virus Species” was identified in the libraries in which the “Virus Species” was found. An “X” indicates that no co-occurrences between these two virus species were found.

### Genetic structure and evolution of Australian sequences

Understanding the evolution of dog viruses is hampered by the lack of sequences available in public databases. Indeed, eight of the 15 dog virus species detected in this study represent the first sequences available from Australia. For many of these species we found multiple distinct sequences within and between sampling locations and, more broadly, Australian states and territories. In some cases, near identical (>99% identity) viruses are detected across states that are separated by over 2,000 km.

Of note, the same viral species were often found in multiple sampling locations and states. For example, CaAstV was present across all three states sampled (Figure 5d). In some cases, these viruses were near identical in sequence. Virus sharing, defined here as sequences sharing >99% pairwise nucleotide identity across the genome, was observed on 13 occasions, 11 of which occurred within sampling locations (11/13). On two occasions near identical virus NoV genotype GVI sequences were detected in Central West and Hope Vale, sharing 99.2% nucleotide identity across the genome (Figure 6b, c). The second was a CanPV sequence that shared 99.8% identity between Central West and Coffs Harbour shelters (Figure 6d). Conversely, we also observed extensive sequence diversity for a given virus species within a sampling location. For example, CaAstV identified in the huntervalley_6 and huntervalley_7 libraries shared only 89% nucleotide identity. On three occasions what appeared to be multiple distinct viruses belonging to the same species were detected within a given library. These included three CaAstVs in hopevale_4, three CaCV in perth_5 and two CPDVs in perth_2 libraries. These libraries were pools comprised of three or more faecal samples.

Finally, we tested whether there was an association between pairwise nucleotide identity and the geographical distance between sampling locations for each virus species (i.e., whether sequences from shelters within the same region, such as NSW, share greater similarity compared to sequences from shelters in different regions, such as NSW and WA). Overall, we observed little correlation between geographical distance and genetic distance (Supplementary Table 7, Supplementary Figure 1). The exceptions were the small positive correlations observed for CanPV (R = 0.78, p = 0.050) and CaAstV (R = 0.67, p = 0.023), indicating that as geographic distance increased, so too did genetic distance.

## Discussion

Viruses cause disease in both domestic and wild dogs (including dingoes) in Australia, resulting in health and financial burdens due to treatment, control, and biosecurity efforts. Given the close contact between dogs, humans, and wildlife, these viruses can have complex ecology with multiple host ranges and spillover events (Day et al. 2012; He et al. 2022; Wu et al. 2024). However, aside from a few well-characterised pathogens like CPV and CHV, genomic data are limited for many of the dog viruses prevalent overseas, restricting understanding of potential population-level impacts and disease ecology that may be relevant to wild and domestic animal management as well as public health.

Australia’s geographic isolation, strict biosecurity measures, and native dingo populations provide a unique opportunity to study viral evolution and transmission. Through a metatranscriptomic approach we characterised the viruses in dogs and dingoes from a variety of geographical locations and sample types, revealing the presence of 86 vertebrate associated viruses belonging to 16 distinct species. These included important canine and potentially zoonotic pathogens, many of which have not previously been sequenced in Australia.

Of note, we identified the first RNA virus – rotavirus A – in dingoes. In contrast to the only previously known dingo virus—a parvovirus in a dog-dingo hybrid that was closely related to domestic dog sequences (Van Arkel et al. 2019)—these RVA fragments are most closely related to human, and bat-associated viruses. This strongly suggests that the dingo RVA resulted from the cross-species transmission, although the exact source species is uncertain. It is notable that the RVA infected dingo lived in proximity to an urban centre and occupied a den site close to a bat colony. While not detected in the RNA of this dingo, preliminary diet analysis of dingo faeces in Myall Lakes National Park have identified some human derived DNA, most likely from the consumption of faeces (Myall Lakes Dingo/Dapin Project Unpublished data). Although domestic dog associated DNA viruses have previously been detected in dingoes (Van Arkel et al. 2019), this represents the first dingo virus that does not appear derived from those found in domestic dogs, and further shows that dogs and dingoes do not necessarily share the same viruses despite their close genetic relationship. Importantly, there was no clear evidence that the RVA fragments were the result of contamination or misidentification of faeces during sampling. The faecal sample was collected in the exact location where a dingo had been displaying behaviours consistent with having just scent marked 21 hours prior. No domestic dogs have previously been detected at that camera site, and dingoes in this area have high (>99%) dingo genetic ancestry (Cairns et al. 2020). Likewise, there was no strong evidence of contamination during extraction or from diet, such as a dingo consuming human or bat faeces, when the reads from the library in question were classified taxonomically. Most mammalian-associated reads from non-dingo organisms were misannotated bacterial reads, with canine-associated contigs comprising the majority of the mammalian-associated subset in this library. Despite this, because only fragments of VP1, NSP1, and NSP3 were identified, further work is needed to recover the complete genome of this virus and confirm its association with dingoes.

We detected six known pathogens associated with canine infectious respiratory disease syndrome (CnPnV, CHV, and CRCoV) and gastroenteritis (CPV, CCoV, and RVA). We also frequently detected CaKoV, CaVV, CaAstV, SaV, NoV, RVI in faecal samples, although they have only limited associations with gastroenteric disease and are largely uncharacterised outside their detection in symptomatic cases (Caddy and Goodfellow 2015; Li et al. 2011; Mihalov-Kovács et al. 2015; Ntafis et al. 2010; Renshaw et al. 2018; Ribeiro et al. 2017).

Despite the presence of six known canine pathogens—some at high abundance—there was strikingly little association between virus detection and clinical disease. It is important to note that there were no reports of diarrhoeal or gastrointestinal disease among the dogs sampled, and only one report of respiratory symptoms (kennel cough), from which CnPnV was recovered. These results suggest that these viruses—including zoonotic viruses like RVA—may cause transient or subclinical infections in dogs yet still be present in high abundance. It is possible that there as unreported diarrhoeal or gastrointestinal disease in these dogs or that disease occurred outside the observational window. This finding highlights the importance of including both healthy and sick dogs in pathogen surveillance and control efforts, as well as understanding that the disease manifestation may be multifactorial and associated with additional stressors.

Rotavirus A can cause severe, sometimes fatal, gastroenteritis in both humans and animals. The G3P[3] genotype is common in dogs globally and has occasionally been identified in acute gastroenteritis cases in humans, including in Australia (Roczo-Farkas et al. 2017). These instances are thought to have been zoonotic transmission events originating in dogs, cats, or ruminants (Khamrin et al. 2006; Theamboonlers et al. 2013). While non-human RVAs such as G3P[3] are present in a small percentage of human gastroenteritis cases, through interspecies transmissions these viruses may contribute to reassortment and vaccine escape (Gentsch et al. 2005; Roczo-Farkas et al. 2017). RVAs with a unique genetic constellation, denoted ‘G3-P[3]-I3-R3-C3-M3-A9-N2-T3-E3-H6’, were discovered in Thailand (Theamboonlers et al. 2013), and the RVAs identified in shelter dogs in this study share the same constellation. Notably, this genotype contains segments that appear to be reassortments from human and feline RVA. Combined with the high viral abundance the detected in dog faeces, these findings highlight the potential risk of zoonotic transmission of these viruses.

For many viruses detected in this study, this is the first description of an Australian variant. These new sequences allow for initial evolutionary analyses, and although the number of sequences from Australian dogs was limited, they revealed that Australian viral lineages form multiple distinct clades within several virus groups (e.g., CanPV, SaV, CaCV, CaKoV, and mamastroviruses). In addition, the number of distinct clusters among Australian sequences suggests that there may have been multiple introductions of these viruses into Australia. However, there was often sequence divergence (>1% across the genome) between Australian sequences and the closest related sequences from overseas, complicating the reconstruction of potential transmission routes into Australia. This is likely due to a combination of older introductions and limited global sampling.

It is tempting to hypothesise how these viruses may have entered Australia. One possible explanation is the importation of domestic dogs from overseas, although introduced feral red foxes (*Vulpes vulpes*) are also common on the mainland. Around 6,000 dogs are imported into Australia annually under strict importation protocols. With the exception of a select few countries (New Zealand, Norfolk Island, and Cocos Island), all are required to stay a minimum of 10 days in quarantine which can extend up to 180 days in some circumstances (https://www.agriculture.gov.au/about/news/on-the-record/importing-dogs-aus). Dogs are required to be vaccinated against Rabies lyssavirus and recommended canine distemper, hepatitis, parvovirus, and parainfluenza vaccination. In combination with additional measures such as veterinary checkups, individual rooms, and strict protocols involving bedding and waste management, this system likely reduces the risk of exotic viruses entering Australia, particularly for known pathogens. Nevertheless, as previously shown for CDV, viral shedding can exceed the current minimum quarantine period, potentially resulting in virus transmission after release (Allen et al. 2023). Several of the enteric viruses detected in this study are environmentally stable for extended periods of time and capable of transmission by fomites (Boone and Gerba 2007). Given this, it may be possible for these viruses to be transmitted through contaminated clothing or shoes. Other routes, such as the illegal entry of dogs or other carnivores on boats and the importation of dog semen, could also introduce viruses (Charles 2022).

Conversely, some virus clades consisted solely of Australian sequences that are relatively divergent from their international counterparts; for example, we detected a novel lineage of CaVV-like viruses which likely represent a novel species. In some cases (e.g., CaKoVs) one clade comprises members from five sampling locations across two states, spanning much of Australia’s east coast (over 2,000 km), and was distinct from non-Australian CaKoVs (95% nucleotide identity with the nearest relatives). This suggests that these lineages may have persisted and spread across Australia for some time. Notably, there was also little geographical pattern observed in the Australian dog viruses described. This lack of pattern extends to genome identity, as geographically closer locations did not consistently contain more similar viruses, with the exceptions of CaKoV and CanPV. This is clearly apparent within the CaAstVs, in which some sequences sampled in different states share greater sequence identity than those within a single sampling location. The lack of geographical structure indicates virus connectivity/transmission between states, perhaps via the human-mediated interstate movement of dogs. It is also interesting to note that all imported dogs are sent to the same quarantine facility in Mickleham, Victoria. Indeed, despite separation by over 2,000 km, the norovirus GVI sequences identified here shared 99.2% nucleotide identity across the genome.

In sum, through our characterisation of viruses in dogs and dingoes across Australia, we sequenced known dog pathogens in the country for the first time and identified the first RNA virus in dingoes which was distinct from viruses associated with domestic dogs. These findings provide insights into the viral diversity in Australian canines, with implications for understanding patterns in viral movement into and within Australia and the potential zoonotic risks associated with canines and dingoes.

## Data availability

All raw data (fastq files) generated for this study are available in the NCBI SRA database under BioProject PRJNA1222450, BioSample accessions SAMN46782038:SAMN46782107, and SRA accessions SRR32313011-SRR32313080. The viral genomes assembled in this study have been submitted to NCBI/GenBank and assigned accession numbers XXX.YYY. The sequences, alignments, and phylogenetic trees generated in this study are available at: https://github.com/JonathonMifsud/Australian-Canine-Virome

## Acknowledgements

We acknowledge the University of Sydney’s high-performance computing cluster Artemis for providing the computing resources used for this study. We thank Gemma Ma, Nicole Balzer, Deanna Douglas, Hannah Dreaver and Kasey Bridge for assistance with sample collection. Thanks to NSW NPWS staff for assistance with sample collection and storage. The Myall Lakes Dingo/Dapin Project acknowledges the Worimi Traditional Owners on who’s land their research takes place.

## Funding

E.C.H. is supported by an National Health and Medical Research Council (NHMRC) Investigator Fellowship (GNT2017197) and by an Australian Research Council (ARC) Discovery Project grant (DP240101313). J.C.O.M. is supported by an Australian Government Research Training Program (RTP) Scholarship.

## Author contributions

J.C.O.M, E.H, and E.C.H conceptualised the study. J.C.O.M, E.H, A.O, B.J.P, J.H, H.F, B.F.A and S.S assisted in sample collection. J.C.O.M and K.V.B performed sample extraction. J.C.O.M. performed the computational analyses. J.C.O.M. and E.C.H. wrote and prepared the original draft. All authors edited and revised the manuscript. E.C.H. funded the project. E.H. and E.C.H. supervised the project.

## Conflict of interest

The authors declare no conflicts of interest.

## Supplementary Figures

**Supplementary Figure 1.** Association between geographical distance and sequence identity across viral species. Each panel represents a distinct virus species as labelled, with points indicating sequence pairs plotted by geographical distance between sampling locations (km) and pairwise nucleotide sequence identity (%). The red dashed line represents the linear regression fit.

## Supplementary Tables

**Supplementary Table 1.** Sample metadata for each library.

**Supplementary Table 2.** Rotavirus A typing results.

**Supplementary Table 3.** Sequence type, trimming methods, and phylogenetic models.

**Supplementary Table 4.** Contig statistics for each library.

**Supplementary Table 5.** Summary information for the viruses identified in this study.

**Supplementary Table 6.** Data underlying the boxplot in Figure 2.

**Supplementary Table 7.** Data underlying the genetic and geographical distances mantel test.

## Notes

### Competing Interest Statement

The authors have declared no competing interest.

