## Supplementary figures and images for "Diverse Viral Pathogens in Australian Canines: Limited Geographic Structure and the First Detection of an RNA Virus in Dingoes"

### Supplementary Figure 1

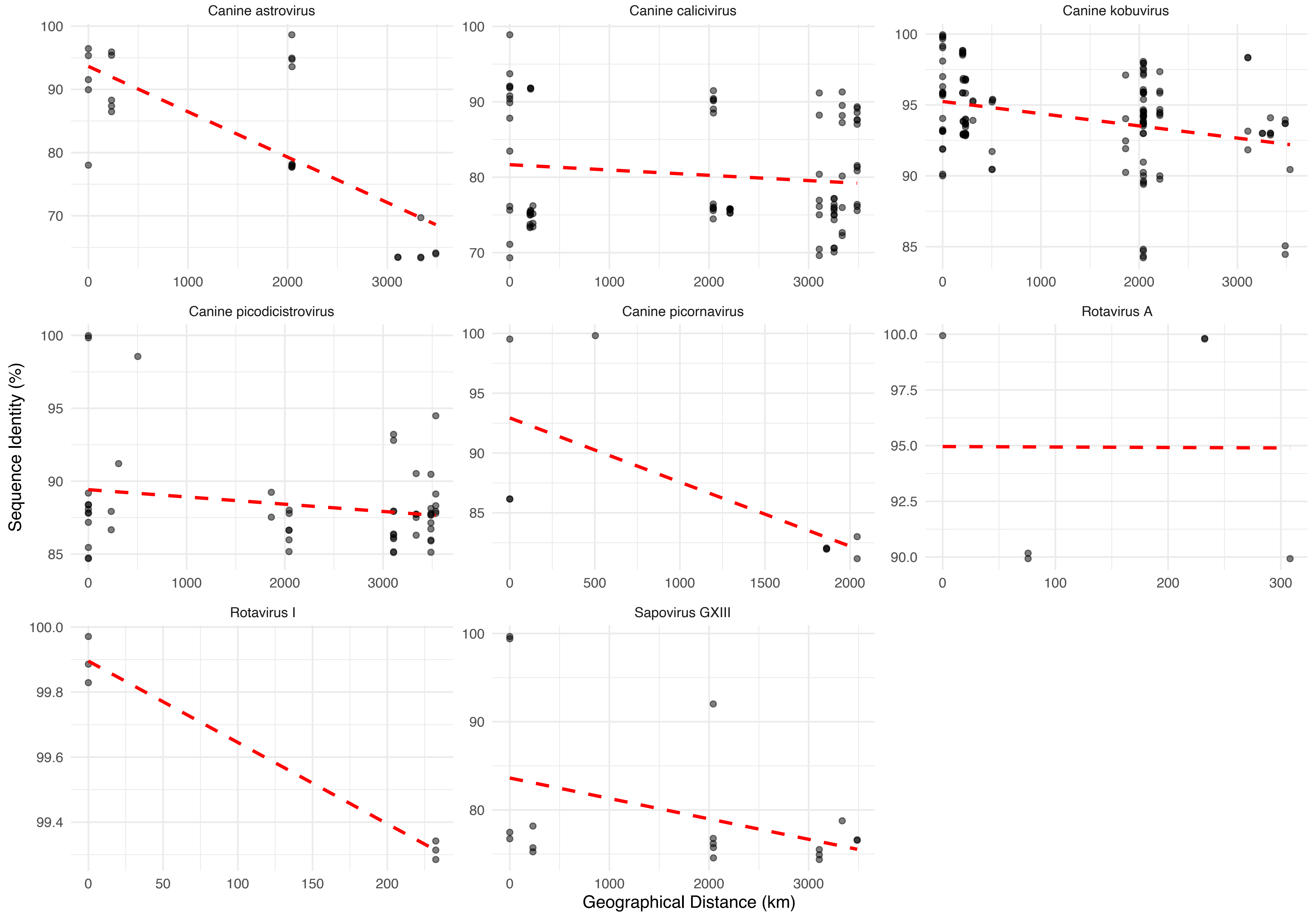
